# The effect of constitutive root isoprene emission on root phenotype and physiology under control and salt stress conditions

**DOI:** 10.1101/2024.02.09.579703

**Authors:** Manuel Bellucci, Mohammad Golam Mostofa, Sarathi M. Weraduwage, Yuan Xu, Mostafa Abdelrahman, Laura De Gara, Francesco Loreto, Thomas D. Sharkey

## Abstract

Isoprene, a volatile hydrocarbon, is typically emitted from the leaves and other aboveground plant organs; isoprene emission from roots is not well studied. Given its well-known function in plant growth and defense aboveground, isoprene may also be involved in shaping root physiology to resist belowground stress. We used isoprene-emitting transgenic lines (IE) and a non-emitting empty vector and/or wild type lines (NE) of *Arabidopsis* to elucidate the roles of isoprene in root physiology and salt stress resistance. We assessed root phenotype and metabolic changes, hormone biosynthesis and signaling, and stress-responses under normal and saline conditions of IE and NE lines. We also analyzed the root transcriptome in the presence or absence of salt stress. IE lines emitted isoprene from roots, which was associated with higher primary root growth, root biomass, and root/shoot biomass ratio under both control and salt stress conditions. Transcriptome data indicated that isoprene altered the expression of key genes involved in hormone metabolism and plant responses to stress factors. Our findings reveal that root constitutive isoprene emission sustains root growth also under salinity by regulating and/or priming hormone biosynthesis and signaling mechanisms, amino acids biosynthesis, and expression of key genes relevant to salt stress defense.

## INTRODUCTION

Isoprene is a non-methane biogenic volatile organic compound (BVOC) emitted by more than half of all terrestrial tropical species, and accounts for the largest flux of BVOCs from the biosphere to the atmosphere (Guenther et al., 2006) and well in excess of anthropogenic hydrocarbon fluxes to the atmosphere. Isoprene is synthesized from the plastidial methylerythritol 4-phosphate (MEP) pathway product, dimethylallyl diphosphate (DMADP) in a reaction catalyzed by isoprene synthase (ISPS). Isoprene emission is metabolically expensive (14 NADPH and 20 ATP/isoprene) for plants (Sharkey and Yeh, 2001), although benefits may outweigh the cost, especially under the conditions like high temperature (Jardine et al., 2012) and oxidative stress (Vickers et al., 2009).

Isoprene is physiologically important to protect the photosynthetic apparatus of chloroplasts from the adverse effects of excessive heat and water-shortage. The underlying proposed mechanisms include stabilization of thylakoid membranes, dissipation of excessive light energy, and protection against oxidative damage by quenching reactive oxygen species (ROS) (Loreto et al., 2001; Velikova et al., 2005; Pollastri et al., 2019). Because ROS accumulation is a common consequence of most abiotic stresses, the antioxidant roles of isoprene underly a general defense mechanism against abiotic stresses (Loreto and Schnitzler, 2010). Recently, the hypothesis of a direct ROS-quenching effect of isoprene has been revised. It is thought that isoprene effects on the transcriptome, proteome, and metabolome accounts for improved stress tolerance (Lantz et al., 2019; Monson et al., 2021; Dani and Loreto, 2022). Indeed, isoprene has a hormone-like activity (Pollastri et al., 2021), acting as a signal molecule for regulating gene expression and biosynthesis and signaling cascades of several plant hormones, including cytokinins (CKs), jasmonic acid (JA), and salicylic acid (SA) (Zuo et al., 2019; Dani et al., 2022; Dani and Loreto 2022; Weraduwage et al., 2023). The control of isoprene over specific metabolite production and defense mechanisms indicates roles of isoprene in growth-defense tradeoffs (Zuo et al., 2019; Monson et al., 2020; Xu et al., 2020; Frank et al., 2021). Isoprene is also known to modulate ROS-mediated cellular signaling for reshaping plant developmental processes (Miloradovic van Doorn et al., 2020).

Isoprene studies have principally focused on the aboveground plant parts (either leaves or whole canopies), while the role of isoprene belowground, especially in the root system, has been rarely investigated. Root isoprene emission was observed in poplar (*Populus x canescens*) (Ghirardo et al., 2011; Miloradovic van Doorn et al., 2020) and in transgenic *Arabidopsis thaliana* harboring the *ISPS* of *Populus x canescens* (*PcIspS,* Loivamäki et al., 2007). The constitutive promoter of *PcIspS* is also active in the roots, particularly in the tips of the fine roots of poplar (Cinege et al., 2009; Miloradovic van Doorn et al., 2020). An upregulation of root growth-related gene *phosphatidylinositol-4-phosphate-5-kinase 2* (*PIP5K2*), *transcription factor* (*MYB59*), and *nitrate transporter* (*NRT1.1*) was found in unstressed isoprene-emitting (IE) leaves of *Arabidopsis* (Zuo et al., 2019). In poplar, the IE line showed a reduction in lateral root (LR) growth but a development of deeper root phenotype when compared with RNAi-lines deficient in isoprene synthesis (Miloradovic van Doorn et al., 2020). These results clearly indicate isoprene connection to plant roots; however, it remains unknown whether plants could use this trait as an advantage to moderate stress resilience as observed in case of other BVOCs emitted from roots (Asensio et al., 2012; Kigathi et al., 2019; Arimura, 2021).

Roots are the anchorage system of plants and are essential for the uptake of mineral nutrients critical for plant growth and productivity. Roots are exposed to several abiotic stresses, such as water-shortage and excess or deficiency of mineral nutrients, and adapt their architecture accordingly (Karlova et al., 2021). Likewise, roots are the primary plant organs that can promptly sense the abnormal accumulation of salts in the soils and initiate signals throughout the plant (Galvan-Ampudia and Testerink, 2011). Salt accumulation is particularly deleterious for plant health in dry climates and is plaguing increasing areas of cultivable lands worldwide because of climate instability (Minhas et al., 2020; Corwin, 2021). In response to salt stress, roots can readjust their growth, dynamics, and architecture and mount defense mechanisms to overcome adverse consequences (Zou et al., 2022). This often requires signaling events to reprogram the architecture through gene-to-metabolite networks, resulting in avoidance and/or heightened protection against salt stress effects (Julkowska et al., 2017).

Recent genetic studies using different plant systems, including tobacco (*Nicotiana tabacum*), poplars (*Populus* spp.), and *Arabidopsis* report compelling evidence for isoprene signaling roles in enhancing plant defense mechanisms (Zuo et al. 2019; Miloradovic van Doorn et al., 2020; Dani et al., 2022). Several studies also indicate that isoprene emission from foliar organs play vital roles in plant physiological responses under various abiotic stresses (Vickers et al., 2009; Loreto and Schnitzler 2010; Jardine et al., 2012). However, little is known about isoprene emission of roots, and its consequences on root physiology. Moreover, it is currently unknown whether root isoprene emission is regulated under any environmental stress, and if root isoprene emission has any effect on root metabolomes and transcriptomes.

In the current study, we used empty-vector (EV) and wild-type (WT) (both non-emitters, NE), and isoprene emitting (IE) transgenic *Arabidopsis* lines (B2 and C4) to test root isoprene effects when exposing plants to salt stress. We investigated physiological responses of roots and the levels of phytohormones and stress metabolites, as well as the changes in MEP pathway metabolites in IE and NE plants under normal and saline conditions. We also examined the transcriptome levels by performing RNA-sequencing (RNA-seq) in the roots of IE and NE lines in presence and absence of salt stress. We show that induction of isoprene emission promotes primary root growth, altering root metabolome (hormone, MEP metabolites, and amino acids) and transcriptome, under control and salt stress conditions.

## RESULTS

### Root isoprene emission, root growth and biomass accumulation

Isoprene-emitting B2 and C4 *A. thaliana* lines developed a deeper root phenotype, showing a higher PR growth under control conditions than the EV NE line (Fig. 1a,b,c,d). This also occurred in soil, where 2-wk-old IE lines showed a 30% higher PR growth with respect to NE (EV) roots (Supplemental Fig. S1).

**Figure 1.**
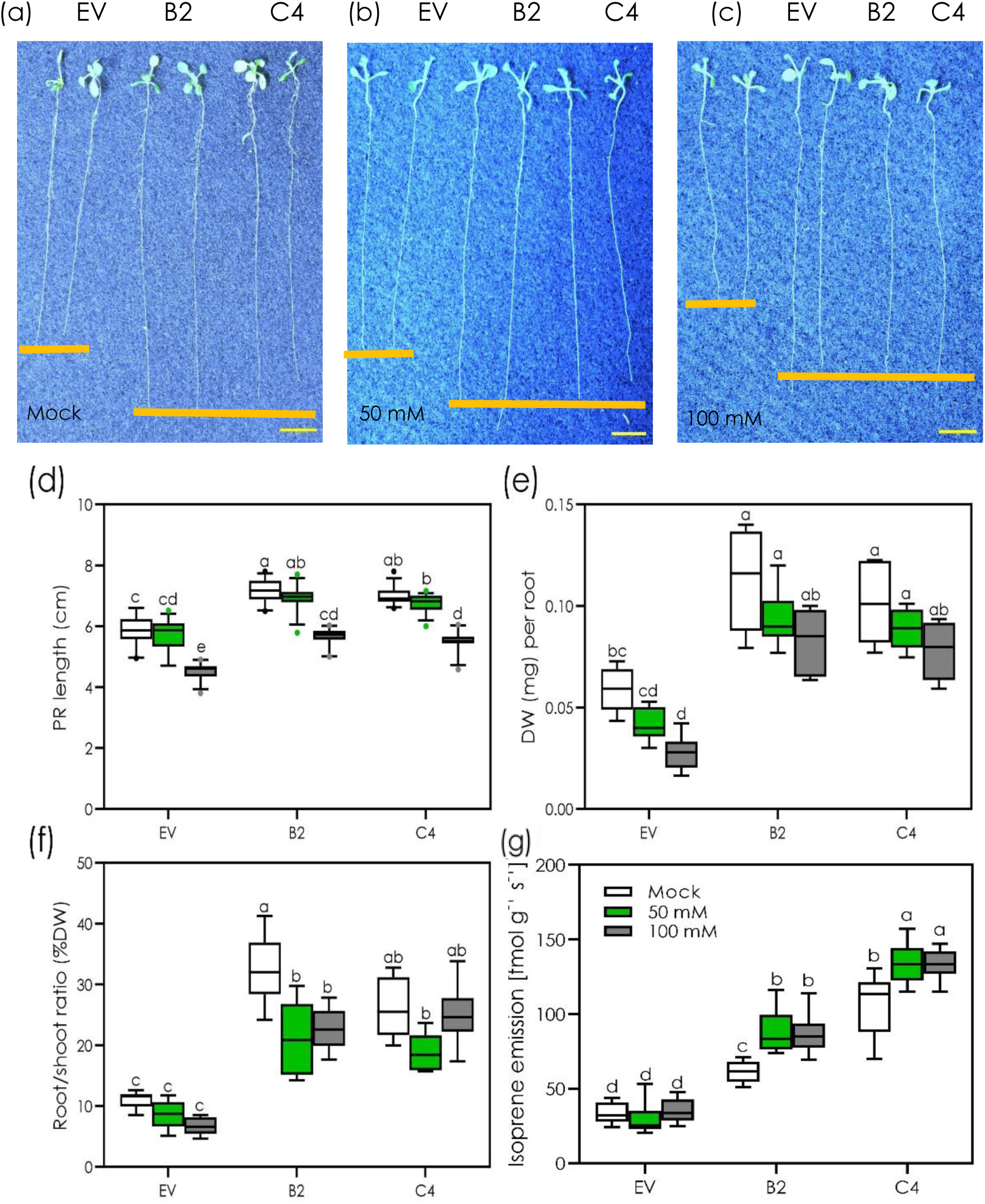
Root phenotypes, growth parameters and isoprene emission of non-emitting (EV) and emitting (B2 and C4) Arabidopsis lines under control and salt-stressed conditions. Representative root phenotype images (a-c), primary root (PR) length (d), dry weight (e), root/shoot ratio (f) and isoprene emission (g) of EV, B2, and C4 were recorded after exposure to 0 (Mock), 50, and 100 mM of NaCl for five days (n=4 plates; 12 seedling per plates). Yellow mark represents scale bar 1 cm. The horizonal line within the box corresponds to the median and the box marks the lower and upper quartiles. All data are means ± standard deviations. Green and grey points indicated outlier data. Different letters above the box plots indicate significant differences (P < 0.05) among the treatments and genotypes, based on two-way ANOVA and Tukey test.

Mild salt stress (50 mM NaCl) caused non-significant reductions of PR length in both NE and IE lines, while severe salt stress (100 mM NaCl) resulted in a 30% and 20% reduction of PR length in NE and IE lines, respectively (Supplemental Fig.S2a).

IE lines showed significantly higher root biomass under both normal and salt stress conditions with respect to NE (Fig. 1e). At 50 mM NaCl, the root biomass of NE and IE lines did not change compared to their respective control conditions (Fig. 1e). Under severe salt stress, root DW was significantly reduced by 60% in the NE line, but only by 30% and 25% in the IE lines B2 and C4, respectively, compared to control conditions (Fig. 1e; Supplemental Fig. S2b). IE lines also showed higher root/shoot ratio than NE, in control and under mild and severe salt stress conditions (Fig. 1f), while no significant changes appeared in the shoot biomass between NE (slightly higher) and IE (Supplemental Fig. S3a).

A measurable level of constitutive isoprene emission was recorded in the roots of the IE lines B2 and C4 lines, whereas isoprene emission was barely detectable in the EV line (Fig. 1g). Salt stress affected the emission of isoprene by increasing its emission by 30% and 20%, respectively in B2 and C4. Constitutive IE did not alter the number of LR in roots of B2 and C4 lines (Fig. S3b).

Isoprene emission affected root gravitropic response in IE lines (Supplemental Fig. S4). When rotating the plates of unstressed five-d-old seedlings by 90°, IE lines responded significantly more than EV line at 2, 5, and 7 h after rotation, which resulted in a steeper phenotype in B2 and C4 lines 24 h after plate rotation (Supplemental Fig. S4).

### Pyruvate and MEP pathway metabolites

Salinity caused a big increase in pyruvate in the EV line. The increase in IE lines was much smaller and was not statistically significant in line C4 (Fig. 2a). IE lines consistently maintained a higher level of MEP pathway metabolites compared to the NE EV line under both control and salt-stressed conditions (Fig. 2b,c,d,e,f,g). In particular, the levels of DXP, MEP, CDP-ME, MEcDP, HMBDP, and IDP+DMADP were roughly two times higher in IE than in NE lines under control conditions. HMBDP decreased in salt-stressed IE lines only, while slightly increasing (15%) in EV (Fig. 2f). IDP+DMADP significantly increased in salt-stressed IE lines compared to control conditions (Fig. 2g).

**Figure 2.**
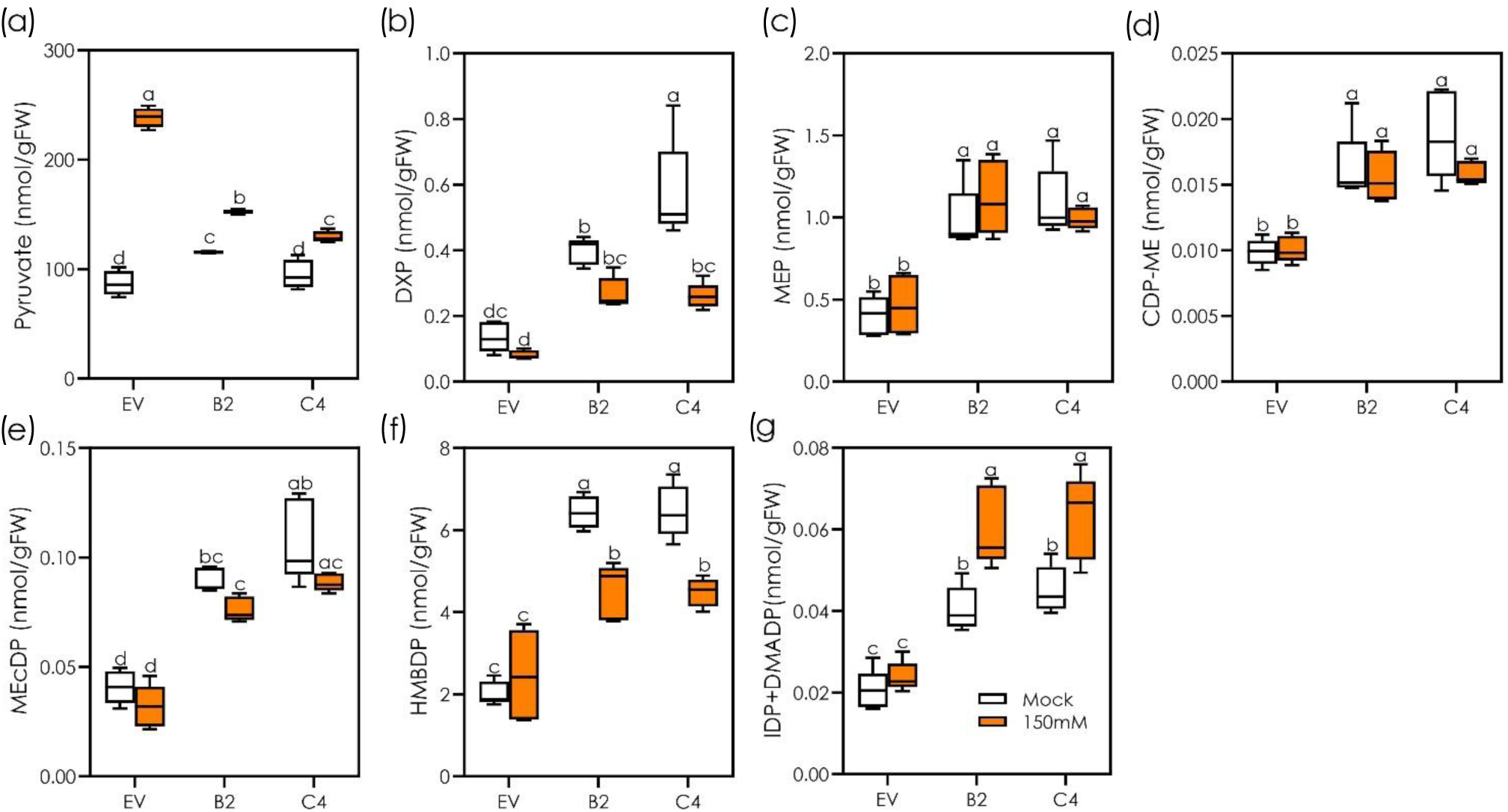
Levels of MEP metabolites in the roots of non-emitting (EV) and emitting (B2 and C4) Arabidopsis lines under control and salt-stressed conditions. Ten-day-old EV, B2, and C4 plants were exposed to 0 (Mock) and 150 mM NaCl solutions for 4 h and the levels of (a) pyruvate, (b) 1-deoxy-d-xylulose 5-phosphate (DXP), (c) 2-C-methylerythritol 4-phosphate (MEP), (d) 4-diphosphocytidyl-2-C-methylerythritol (CDP-ME), (e) 2-C-methyl-D-erythritol 2,4-cyclodiphosphate (MEcDP), (f) (E)-4-Hydroxy-3-methyl-but-2-enyl diphosphate (HMBDP), (g) isopentenyl diphosphate (IDP) and dimethylallyl diphosphate (DMADP) were recorded (*n*=4). The horizonal line within the box corresponds to the median and the box marks the lower and upper quartiles. All data are means *±* standard deviations. Different letters above the box plots indicate significant differences (P < 0.05) among the treatments and genotypes, based on two-way ANOVA and Tukey test.

### Hormones

When compared to the NE line, IE lines showed a significant increase in the levels of OPDA, JA, and JA-Ile while exhibiting a similar level of MeJA under control conditions (Fig. 3a, b, c, d). In response to salt stress, NE displayed a significant increase in the levels of OPDA, JA, and MeJA but a non-significant change in JA-Ile level when compared with control conditions (Fig. 3a, b, c, d). On the other hand, the levels of JA, OPDA, and MeJA but not JA-Ile significantly decreased in salt-stressed IE B2 and C4 lines relative to control conditions (Fig. 3a, b, c, d). The levels of ABA were constitutively very low in both NE and IE lines under control conditions (Fig. 3e). Salt stress resulted in a significant increase in the ABA level by 30, 20 and 25 times in EV, B2, and C4 lines, respectively, in comparison with control conditions.

**Figure 3.**
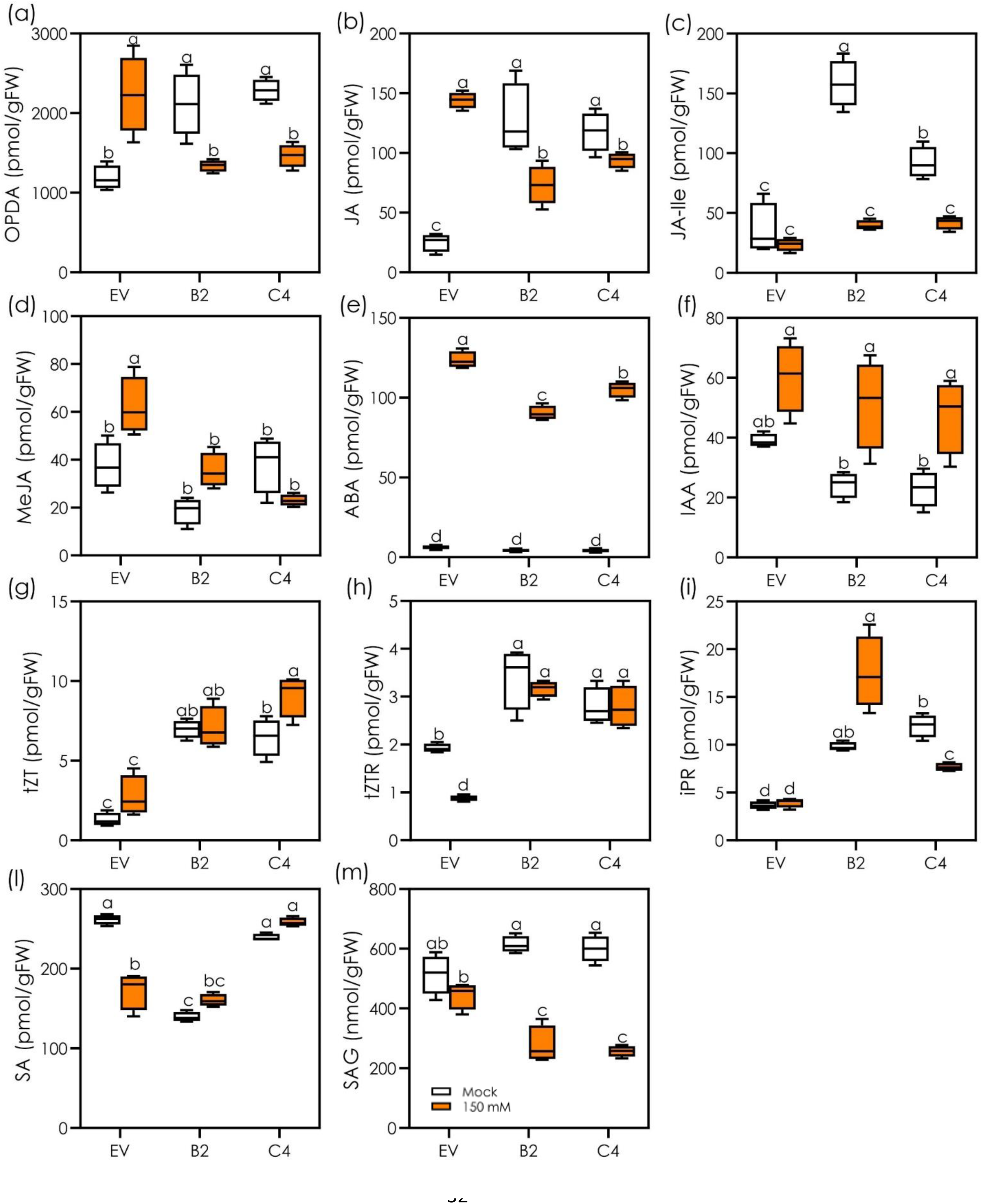
Effects of root isoprene emission on hormone levels in the roots of non-emitting (EV) and emitting (B2 and C4) Arabidopsis lines under control and salt-stressed conditions. The levels of different hormones and hormonal precursors, including (a) oxylipin 12-oxo-phytodienoic acid (OPDA), (b) jasmonic acid (JA), (c) JA-isoleucine (JA-Ile), (d) methyl-jasmonate (MeJA), (e) abscisic acid (ABA), (f) indole acetic acid (IAA), (g) trans-zeatin (tZT), (h) trans-zeatin riboside (tZR), (i) isopentenyl riboside (iPR), (l) salicylic acid (SA), and (m) salicylic acid glucoside (SAG) were determined in root tissues of EV, B2, and C4 lines after exposure to 0 (Mock) and 150 mM NaCl solution for 4 h (*n*=4). The horizonal line within the box corresponds to the median and the box marks the lower and upper quartiles. All data are means ± standard deviations. Different letters above the box plots indicate the significance differences (P < 0.05) among the treatments and genotypes, based on two-way ANOVA and Tukey test.

IAA levels did not differ significantly among the lines in control conditions (Fig. 3f). However, IAA levels increased significantly in salt-stressed IE lines but to a non-significant degree in the NE line (Fig. 3f). IE lines maintained a constantly higher level of cytokinins, including tZ, tZR, and iPR, than the NE line under both control and salt stress conditions (Fig. 3g, h, i). In the presence of salt stress, tZR level significantly decreased (Fig. 3h) whereas the levels of tZ and iPR remained at the control level in NE plants (Fig. 3g,i). On the other hand, all cytokinins remained significantly elevated in IE lines under salt stress (Fig. 3g,h,i).

Without salt stress, SA level was comparable in EV and C4 lines, but significantly lower in the IE B2 line (Fig. 3l). Salt stress severely decreased the level of SA (by 35%) in the NE line but had no significant effect on the level of SA in IE lines (Fig. 3l). The level of SAG was comparable among the lines under control conditions (Fig. 3m). Salt treatment caused a significant decline of SAG in IE lines but not in the NE line when compared with control conditions (Fig. 3m).

### Amino acids and organic acids

The constitutive isoprene emission did not generally affect the levels of amino acids, including asparagine, glutamate, and aspartate, whereas a lower level of proline was recorded in IE lines when compared with EV line (Fig. 4a, b, c, d). In the presence of severe salinity, EV line showed a significant increase in asparagine level and decline in glutamate level, while proline and aspartate levels were not statistically different in comparison with control conditions (Fig. 4a, b, c, d). Salt exposure resulted in a remarkable increase in the levels of asparagine, proline, glutamate, and aspartate in B2 and C4 lines (Fig. 4a, b, c, d). Organic acids, including citrate, succinate, fumarate, malate, glycerate, and glycolate showed differential responses in both EV and IE lines under normal and salt stress conditions (Supplemental Fig. S5). The levels of citrate, succinate, and fumarate were significantly higher in EV line when compared with IE lines under salt-unstressed conditions (Supplemental Fig. S5a, b, c). On the other hand, malate and glycolate levels were significantly higher in IE line B2 in comparison with EV line under control conditions (Supplemental Fig. S5d, e). Salt stress led to a significant decrease of citrate, succinate, and fumarate levels, while malate and glycolate levels significantly increased in EV line when compared with control conditions (Supplemental Fig. S5a, b, c, d, f). In IE lines, salt exposure resulted in a significant increase in citrate level in C4 line and glycerate and glycolate levels in B2 line relative to those levels found in control conditions (Supplemental Fig. S5a, e, f). The levels of succinate, fumarate, and malate did not show significant alteration in IE in response to salt stress when compared with control conditions (Supplemental Fig. S5b, c, d).

**Figure 4.**
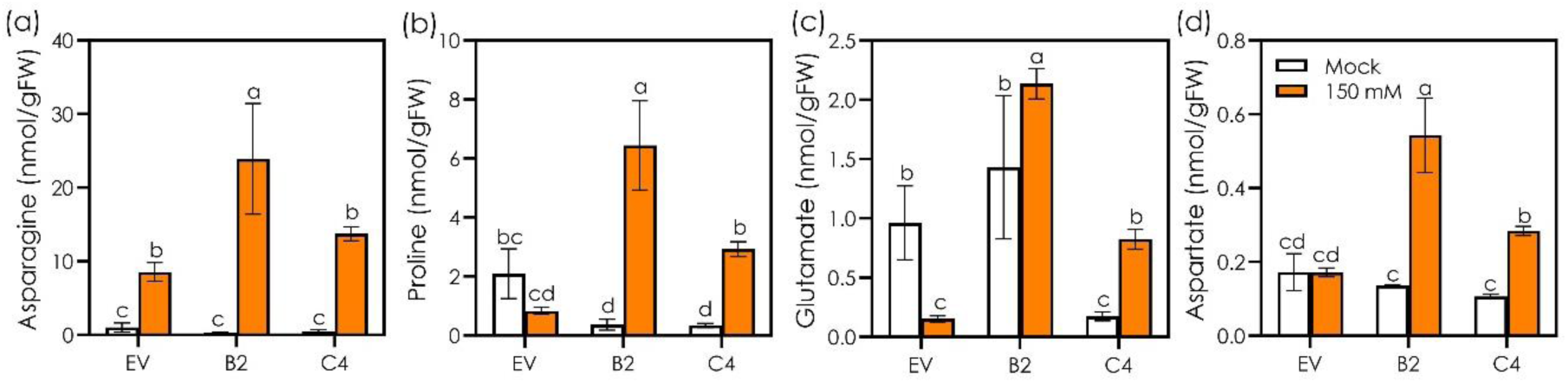
Effects of root isoprene emission on the levels of stress-associated amino acids in the roots of non-emitting (EV) and emitting (B2 and C4) Arabidopsis lines under control and salt-stressed conditions. The levels of asparagine (a), proline (b), glutamate (c), and aspartate (d) were measured in root tissues of EV, B2, and C4 plants after exposure to 0 (Mock) and 150 mM NaCl solutions for 4 h. All data are means ± standard deviations of 4 biological replicates. Different letters above the box plots indicate significant differences (P < 0.05) among the treatments and genotypes, based on two-way ANOVA and Tukey test.

### Root transcriptome profiles

Transcriptome analysis was conducted on 10-d-old roots of WT and EV (NE lines) and B2 and C4 (IE lines) after a four h exposure to 0 mM and 150 mM NaCl. The transcriptome profiles of NE and IE genotypes were analyzed using principal component analysis (PCA) and t-distributed stochastic neighbor embedding (t-SNE) statistical methods to identify the transcriptomic changes induced by IE capacity, salt stress or their interaction, as depicted in Fig. 5a, b. A high proportion (52%) of the variance in the transcriptome changes was made by the effect of salt and was captured by PC1, whereas a low proportion (10%) of variance was captured by PC2, representing the effect of genotypes (Fig.5a). Both PCA and t-SNE separated the investigated genotypes exposed to 0 mM and 150 mM NaCl into main clusters (Fig. 5a, b). Notably, the predominant transcriptome variations across the investigated genotypes were due to salt stress, as denoted by the larger variances in PC1 and the t-SNE Y-axis (Fig. 5a, b). On the other hand, minor transcriptome changes were attributed to genotype-specific effects as indicated by PC2 and t-SNE X-axis (Fig. 5a, b).

**Figure 5.**
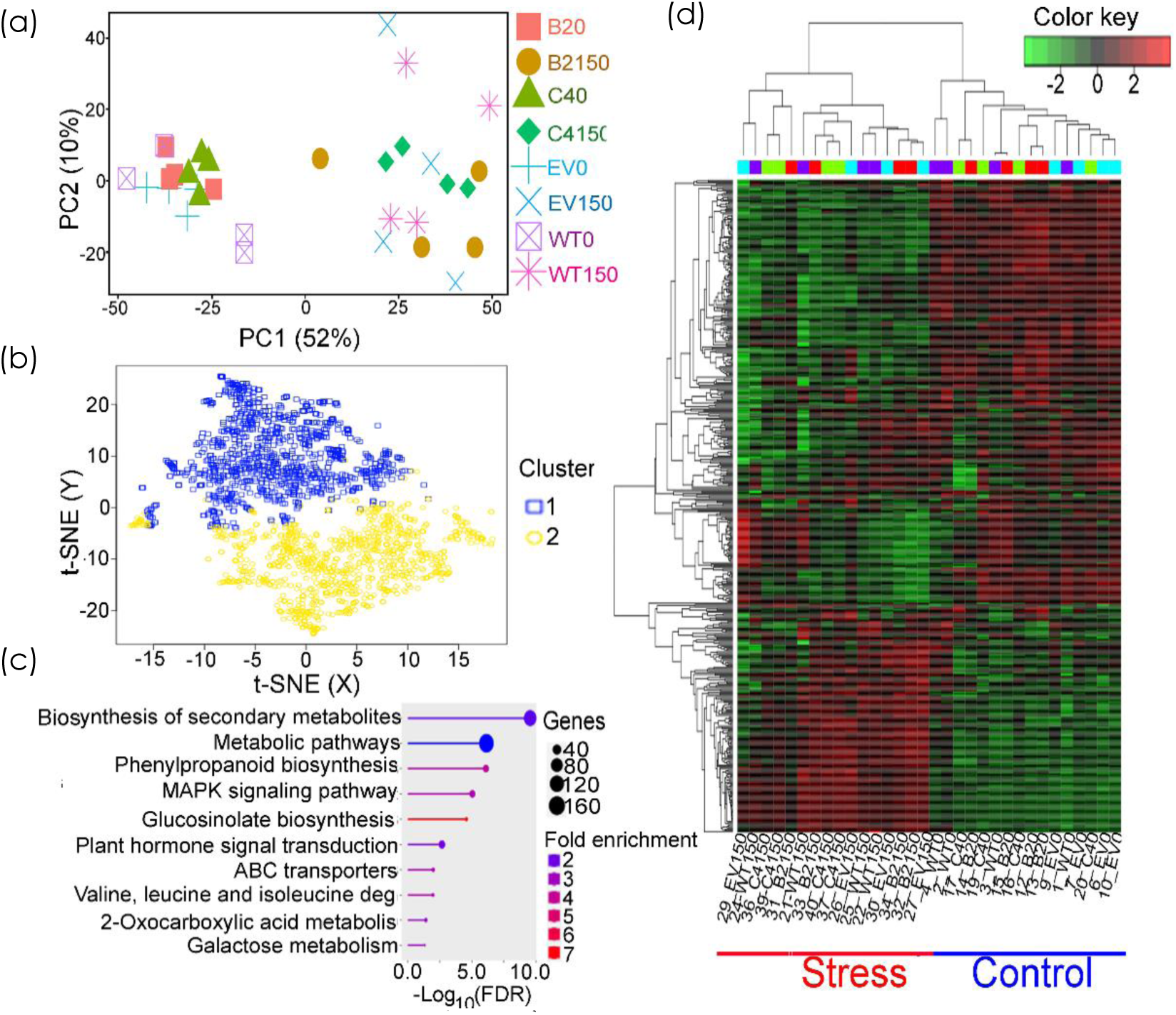
The transcriptome profiles of isoprene non-emitter wild-type (WT) and empty vector (EV) plants, alongside isoprene emitter transgenic lines B2 and C4 in response to 4 hours 0 mM NaCl (WT0, EV0, B20 and C40, respectively) and 150 mM NaCl salinity stress (WT150, EV150, B2150 and C4150, respectively) conditions. (a-b) Principal component analysis (PCA) and t-distributed stochastic neighbor embedding (t-SNE) plots of the transcriptome changes in the investigated genotypes in response to salinity stress. (c) KEGG enrichments analysis of the top 500 variable genes associated with salinity stress responses in the investigated genotypes. (d) Heatmap hierarchical clustering of the normalized expression top 100 variable genes in the investigated genotypes in response to stress and control conditions. Color bar indicates high (red) and low expression (green) levels for each genotype under different treatment conditions.

To get in-depth insights into the key metabolic pathway(s) interlinked with the clustering in PCA and t-SNE for the investigated genotypes, we executed a KEGG pathway enrichment assessment of the transcriptome data using iDEP 0.96 (www.bioinformatics.sdstate.edu/idep96/). This KEGG enrichment revealed that the most enriched genes associated with the genotype separation in PCA and t-SNE belonged to the ‘biosynthesis of secondary metabolites’ and ‘metabolic pathways’ metabolic pathways (Fig. 5c). A significant number of these variable genes also aligned with metabolic pathways tied to stress response mechanisms, such as ‘phenylpropanoid biosynthesis’, ‘MAPK signaling’, ‘glucosinolates biosynthesis’, and the ‘plant hormone signal transduction pathway’. The heatmap hierarchical clustering, presented in Fig. 5d, shows the top 100 variable genes across genotypes and salt stress conditions, distinctly emphasizing the pronounced impact of salinity stress.

### Transcriptome analysis, gene ontology, KEGG enrichment, and protein-protein network

Log_2_ FC ≥ 1 (up-regulated) and Log_2_ FC ≤ −1 (down-regulated) with FDR ≤ 0.05 were used as minimum cutoffs to identify DEGs that are robustly regulated by isoprene emission capacity, salinity stress, or both. The DEG analysis revealed that 863 and 1010 genes were up-regulated, whereas 756 and 577 genes were down-regulated when comparing salt-stressed and control NE (WT and EV, respectively) (Fig. 6a). Likewise, 1007 and 919 genes were up-regulated, whereas 716 and 666 genes were down-regulated when comparing salt-stressed and control IE (B2 and C4, respectively) (Fig. 6a). The constructed Venn diagram illustrated that there were 733 overlapping DEGs (513 up-regulated and 220 down-regulated) when comparing all IE and NE genotypes under control and salt stress conditions (Fig. 6b). These represent the core transcriptome changes in response to salinity stress regardless of isoprene emission capacity. In contrast, 373 up-regulated genes and 415 down-regulated genes were identified only in IE lines under control and salt stress conditions (Fig. 6b). These 788 overlapping DEGs represent core specific transcriptome changes modulated by the presence of isoprene, which may contribute to change of root physiology in presence of salinity stress (Fig. 6b).

**Figure 6.**
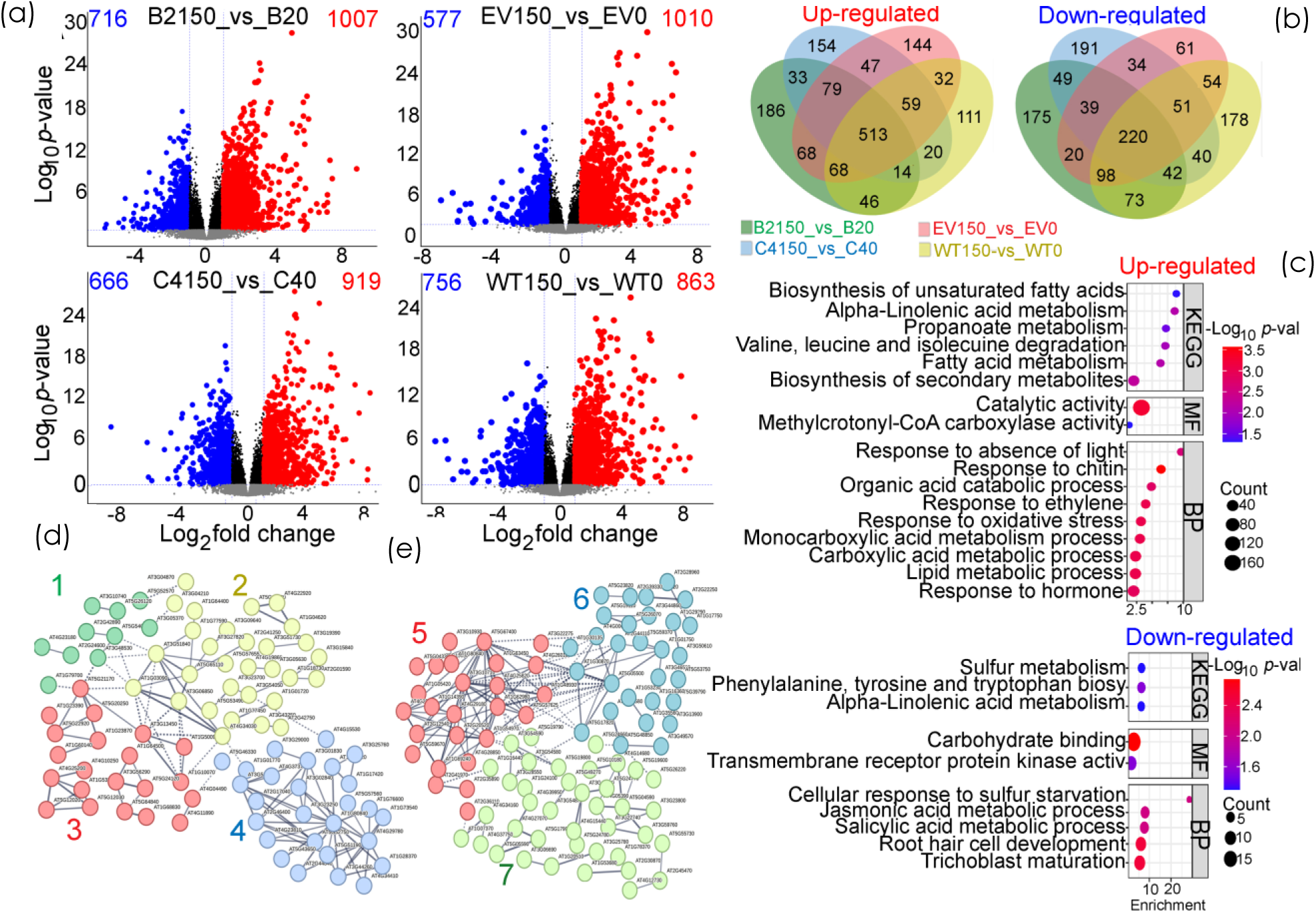
Differential expressed genes (DEGs) of isoprene non-emitter wild-type (WT) and empty vector (EV), and isoprene emitter transgenic lines B2 and C4 in response to 4h 0 mM NaCl (WT0, EV0, B20 and C40, respectively) and 150 mM NaCl (WT150, EV150, B2150 and C4150, respectively). (a) Volcano plots of significantly up-regulated [log_2_ (fold-changes) ≥ 1; *q*-values ≤ 0.05], and down-regulated [log_2_ (fold-changes) ≤ −1; *q*-values ≤ 0.05] genes in ‘WT150 vs. WT0’, ‘EV150 vs.EV0’, ‘B2150 vs. B20’ and ‘C4150 vs. C40’ comparisons. Blue-dashed lines represent the q-value and fold-change threshold. Red and blue points highlight the up-regulated and down-regulated genes, respectively, in the investigated comparisons. (b) Venn diagrams of overlapping DEGs in ‘WT150 vs. WT0’, ‘EV150 vs.EV0’, ‘B2150 vs. B20’ and ‘C4150 vs. C40’ comparisons. Four independent biological replicates (*n* = 4) were collected from each genotype under different treatments for the transcriptome analysis. (c) Gene ontology (GO) and KEGG enrichments analyses of isoprene emitter overlapping DEGs. (d-e) Protein-protein interaction networks of up-regulated (d) and down-regulated (e) overlapping genes in isoprene emitter transgenic B2 and C4 lines in response to salinity stress.

Next, we conducted gene ontology (GO) and KEGG enrichment analysis to get further insight into biological attributes of the 788 overlapping DEGs in the IE B2 and C4 lines (Fig. 6c). The GO enrichment analysis using Fisher’s test and Bonferroni corrections *q*-values ≤ 0.05 classified the 788 overlapping DEGs into two GO categories: (1) molecular functions (MF) and (2) biological processes (BP) (Fig. 6c). The overlapping up-regulated genes in the MF category had highly enriched GO terms related to ‘catalase activity; and ‘methylcrotonyl-CoA carboxylase activity’, whereas ‘carbohydrate binding’ and ‘transmembrane receptor protein kinase activity’ were highly enriched GO terms among down-regulated genes of the MF category (Fig. 6c). Top enriched GO term (fold >2) of the upregulated overlapping DEGs in the MF category was the catalytic activity. In the BP category, highly enriched GO terms showing up-regulated genes were ‘response to absence of light’, ‘response to chitin’, ‘response to organic acid catabolic process’, ‘response to ethylene’ as well as ‘response to oxidative stress’ (Fig. 6c). On the other hand, ‘cellular response to sulfur starvation’, ‘jasmonic acid metabolic process’, ‘salicylic acid metabolic process’ and ‘root hair cell development’ were those with GO enriched terms in down-regulated (Fig. 6c).

With respect to KEGG enrichments, ‘biosynthesis of unsaturated fatty acids’, ‘propanoate metabolism’, ‘valine, leucine and isoleucine degradation’ and ‘fatty acid metabolism’, and ‘biosynthesis of secondary metabolites’ were among the enriched metabolic pathways in the up-regulated overlapping genes (Fig. 6c). On the other hand, ‘sulfur metabolism’ and ‘phenylalanine, tyrosine and tryptophan biosynthesis’ were among the enriched KEGG pathways in the down-regulated overlapping genes (Fig. 6c).

Finally, a protein-protein interaction (PPI) networks analysis for the target overlapping DEGs associated with IE B2 and C4 lines in response to salinity stress was performed (Fig. 6d). PPI networks demonstrated that ‘(1) auxin biosynthesis’, ‘(2) fatty acid biosynthesis and metabolism, citrate cycle, and valine, leucine and isoleucine biosynthesis and degradation’, ‘(3) small heat shock protein, suberin biosynthesis and B-box zinc finger’, ‘(4) response to chitin, calmodulin binding protein, and ethylene-activated signaling pathway’ were clustered in the up-regulated overlapping genes in IE lines (Fig. 6d). Whereas PPI networks of ‘(5) phenylpropanoid biosynthesis and secretory peroxidase’ and ‘(6) jasmonic acid biosynthesis process, tryptophan biosynthesis, sulfur metabolism’ pathways were clustered in the down-regulated overlapping genes in IE lines (Fig. 6e).

## DISCUSSION

While the functions of leaf-emissions of isoprene in plant interactions with atmospheric components is well-studied (Loreto and Schnitzler, 2010; Bellucci et al., 2023), the belowground role of isoprene is yet to be clarified, particularly under stress situations. Here, we demonstrate that the capacity to emit isoprene radically altered the phenotypes of the roots stimulating root growth of IE lines under control conditions on 10-d-old Arabidopsis plants grown in artificial substrate (Fig. 1d) and in 2-w-old Arabidopsis plants grown on soil (Supplemental Fig. S1). These isoprene emitting lines invested more in roots than NE plants (Fig. 1f), showing a distinct deeper root phenotype, faster PR growth, and higher biomass (DW basis) both under control and salt-stress conditions (Fig. 1d; Supplemental Fig. S6).

Transgenic *N. tabacum* plants ectopically overexpressing eucalyptus *ISPS* exhibited a dwarf phenotype when compared with the EV’s aboveground growth (Zuo et al., 2019). However, suppression of isoprene emission capacity in poplar resulted in slower growth and reduced apical dominance (Dani et al., 2022). It is likely that natural emitters behaved differently from ISPS-overexpressing lines in resource allocation in the aboveground. Nonetheless, our results suggest that continuous isoprene emission might contribute to the shifting of resources from aboveground to belowground, resulting in a robust root growth and better salt tolerance capacity in IE lines.

There is evidence that the roots of poplar (Ghirardo et al., 2011) and ISPS-overexpressing Arabidopsis (Loivamäki et al., 2007; Miloradovic van Doorn et al., 2020) emit a trace amount of isoprene. We also observed significant root isoprene emission from both IE transgenic lines compared to the EV line under control conditions (Fig. 1g), although, from isoprene concentration in the head space of *A.thaliana* IE roots, a much lower emission than from leaves can be estimated, and a concentration similar to hormones (Dani et al., 2022). The non-negligible emission of isoprene seen in EV roots may reflect non-enzymatic isoprene formation.

Salt stress further increased the level of isoprene significantly in IE roots (Fig. 1g). However, we must consider that VOC diffusion out of roots may be more restricted than in leaves where stomata are present, and that in the plates, and especially in soil, isoprene cannot dissipate as quickly as that emitted from leaves in air. Thus the smaller emission rate might still be associated with significant effects.

It is known that isoprene emission by leaves is sustained under salt stress (Loreto and Delfine, 2000). Our results suggest a stress-responsive role of isoprene in roots, as previously observed in the leaves, especially under heat and drought stress scenario (Velikova and Loreto 2005; Brilli et al., 2007; Jardine et al., 2012; Tattini et al., 2014).

Isoprene synthases are located in plastids (Sharkey et al., 2013), however, recently, Zhou and Pichersky (2020) reported that a TPS-b type TPS47 is localized in the cytosol of tomato (*Solanum lycopersicum*). *TPS47* encoded a functional ISPS that can catalyze the formation of isoprene from DMADP *in vitro*. Interestingly, *TPS47* expression was identified in multiple tissues, including root tissues of tomato (Zhou and Pichersky, 2020). We infer that root isoprene emission is plausibly a common trait of the plants that harbor the ISPS genes in roots. However, the DMADP pool is very low in the cytosol (Weise et al., 2013), which perhaps explains why only trace amount of isoprene is emitted by roots in comparison to leaves.

Root isoprene emission in the absence of light may be different from the photosynthesis (light)-dependent isoprene synthesis via the MEP pathway (Sharkey and Yeh, 2001). In photosynthetic organisms, photosynthesis-independent isoprene emission at a rate similar to that emitted photosynthetically was so far identified only in microalgae growing heterotrophically on a sugar-rich substrate (Dani et al., 2020). We showed that root isoprene emission (Fig. 1g) was accompanied by a general stimulation of the levels of pyruvate and the metabolites of the MEP pathway in the root tissues of IE lines (Fig. 2). This suggests that constitutive expression of *EgISPS* resulted in an increased flux of MEP pathway metabolites in the belowground organs of IE lines. Furthermore, salt-induced enhancement of the levels of DMADP and IDP corresponded with an increased emission of isoprene from the roots of both transgenic lines. Many of the MEP pathway enzymes are light-dependent (Sharkey and Yeh, 2001) and isoprene emission from leaves has even been modeled based on light-dependent generation of reducing power (Morfopoulos et al., 2014). Thus, the capacity of roots to emit isoprene may be based on a specific model which is different from that of the leaves. Indeed, our KEGG pathway analysis also revealed an upregulation of DXS in IE roots under salt stress (Supplemental Fig. S7a), suggesting a MEP pathway activation at the transcript level in the roots. DXS catalyzes the first committed step of MEP pathway, which is known to be crucial in maintaining a continuous flux of MEP pathway metabolites (Banerjee and Sharkey, 2014).

The higher PR length was correlated with the enhanced biosynthesis of growth hormones in the roots of IE lines. In comparison with the NE EV, IE lines maintained a consistently high level of CKs (tZ, tZR, and iPR) in their roots under both normal and saline conditions. Dani et al. (2022) also reported an increased level of CKs (iP and iPR) in the leaves of IE poplars compared to transgenic NE plants. KEGG analysis revealed an upregulation of zeatin synthesis in IE lines under control condition (Supplemental Fig. S8a). We propose that isoprene-induced high level of CKs has significant effect on faster growth of PR, and thus played beneficial roles in short-term salinity tolerance. We also observed an enhanced level of IAA in the roots of IE transgenic lines under salt stress conditions. These results suggest that isoprene might have interfered with the CK-auxin crosstalk in maintaining root meristem size, ensuring root growth (Moubayidin et al., 2009), and controlling root development (Dello Ioio et al., 2008).

Interestingly, the levels of JA-associated metabolites like OPDA, JA, and JA-Ile and SA conjugate (SAG) were significantly decreased while the ABA level markedly increased in both IE lines upon exposure to salt stress. KEGG analyses showed an upregulation of several components of JA synthesis in unstressed IE roots (Supplemental Fig. S8b), and this supported the high JA levels detected in unstressed IE roots. High levels of JA-related metabolites (OPDA, JA, JA-ile) in unstressed IE roots, suggest a defense priming effect of isoprene as already demonstrated in leaves (Zuo et al., 2019; Monson et al., 2021).Thus, isoprene might have stimulated a general activation of stress hormones (e.g., ABA) but a lower activation of induced stress responses involving Systemic Induced Resistance (by JA) or Systemic Acquired Resistance (by SA) (Saxena et al., 2020).

Metabolomic analysis also revealed salt-induced upregulation of the stress metabolites like proline, aspartate, and glutamate, as well as the amino acid asparagine. Amino acids, including proline, aspartate, and glutamate are well known salt-tolerant metabolites, acting as osmolytes to support plants under salt-induced osmotic stress (Rai, 2002). Additionally, KEGG analysis showed an upregulation of plant hormone signal transduction pathways mediated by zeatin, ABA, auxin, and JA in the IE roots (Supplemental Fig. S7b), which might be crucial for persistent growth of PR when exposed to salt stress (Fig. 1a).

Acquisition of the capacity to emit isoprene may alter gene expression aboveground (Behnke et al., 2010), and the possibility that isoprene acts directly as a signaling molecule was proposed by Harvey and Sharkey (2016). Previous transcriptomic analysis of unstressed isoprene-fumigated Arabidopsis leaves showed that isoprene acts as a signaling molecule upstream of many growth regulators involved in various signaling cascades (Zuo et al., 2019). In isoprene-treated Arabidopsis leaves, there was an upregulation of JA signaling/biosynthesis genes (Zuo et al., 2019), whereas in IE roots, we observed a downregulation of *ROXY20*, which negatively regulates JA signaling and a wounding-inducible JA-related gene (*CYP84A4*) (Table S2). Moreover, we observed an upregulation of ABA-associated genes (*CYP709B2, ARCK1, and BBD2*). Interestingly, isoprene-fumigated Arabidopsis leaves showed a downregulation of several ABA-related genes (Zuo et al., 2019). For instance, the *bifunctional nuclease in basal defense response 2 (BBD2)*, which is involved in JA and ABA signaling for drought tolerance (Huque et al., 2021), was downregulated (Zuo et al., 2019) while it was upregulated in salt-stressed roots of IE lines (Table S2). Root IE capacity upregulated the expression of genes associated with gibberellin deactivation (*ATGA2OX1*), brassinosteroid signaling (*JMJD5*), but downregulated *methyl esterase 1* (*MES1*), which is known to encode MES1 that plays a role in methylsalycylate (MeSA) or methyl indole-3-acetic acid (Me-IAA) hydrolysis (Vlot et al., 2008; Yang et al., 2008) (Table S2). Exogenous Me-IAA (the inactive form of IAA) was found to significantly inhibit root growth in plant (Yang et al., 2008).

In the IE B2 and C4 lines, we also found an enhanced expression of several key genes associated with the defense against biotic and abiotic stresses, as also reported by Zuo et al. (2019) in isoprene-treated Arabidopsis leaves. Particularly, we observed an alteration of associated genes involved in general defense responses (*G1IP-LIKE, PTF1, SPA2, ROXY20, NF-YC13*), in biotic stress responses (*LOV1, MAGL6,* and *CYP709B2*) and in abiotic stress response, like salt stress (*ARCK1* and *CYP709B2*), light (*HYR1* and *OEP6*), temperature (*JMJD5*), oxidative (*MDAR4* and *ELT3*), osmotic (*ARCK1, CYP709B2, MAGL6, NAC080*, and *CYP84A4*) and drought (*BBD2*) (Table S3). Finally, protein networks analysis revealed an upregulation of small heat shock protein and calmodulin binding proteins in IE lines, which perhaps contributed to enhanced salt tolerance of IE lines.

## CONCLUSION

A tiny emission of isoprene from roots can be measured; given reduced diffusion through soil this could have physiological significance. A high investment of biomass into roots was observed as a consequence of constitutively expressed root isoprene emission. Possible causes include i) hormone stimulation, including ABA, auxin, and CK; ii) improved synthesis of stress tolerance metabolites like proline, aspartate, and glutamate; and iii) modulation of the hormone signaling pathways (e.g., JA, SA, CKs, and ABA) and associated gene expression. These findings suggest the onset of pleiotropic effects when isoprene is emitted, affecting gene expression, hormone synthesis, and the phenotypes of plants. In particular, the belowground signaling role of isoprene modulating hormones and protecting roots from stresses opens up a promising avenue for future research on isoprene. In accordance with Sharkey & Monson (2017), our findings imply that isoprene has a general role in regulating biological processes that are critical for conferring plant stress resilience. Further comparative studies of transgenic lines that are capable or defective in isoprene emission across diverse plant species will provide deeper insights into the specific mechanisms by which isoprene alters root physiology to improves plant resistance to salinity and other soil-borne stresses.

## MATERIAL AND METHODS

### Plant materials, growth conditions, and salt treatments

We used our previously developed *A. thaliana ISPS*-transgenic lines (Zuo et al., 2019). The full complementary *ISPS* DNA sequence including native transit peptide sequence from *Eucalyptus globulus* was placed under the *Arabidopsis* rubisco small subunit promoter rbcS-1A (*At1g67090*) to generate the transgenic, IE lines. After the *ISPS* sequence, an *octopine synthase* gene was placed as a transcriptional terminator of the transgene. The construct without *EgISPS* was also transformed with the vector to generate a NE EV control. After successful transformation, seven transgenic lines and three empty vector lines were characterized. Among the transgenic lines, B2 and C4 showed highest level of isoprene emission (Zuo et al., 2019). Thus, we selected B2 and C4 transgenic lines as IE, together with EV-B3 (referred as EV hereafter) as NE for the current study.

The *A. thaliana* seeds were surface sterilized with 70% ethanol and 5% bleach solution for 5 and 10 min, respectively, followed by washing with sterilized milli-Q water for five times. Seeds were placed on germination medium (GM) plates containing ½ Murashige and Skoog (MS) solid medium (0.8% agar and 1% sucrose), supplemented with Gamborg’s vitamins solution (Sigma-Aldrich, Germany). After stratification for three days at 4°C in the dark, the plates were vertically placed in the growth chamber under a 16 h: 8 h, light: dark photoperiod, 120 μmol m^−2^ s^−1^ photosynthetically active radiation (PAR), 23/20°C day/night temperatures, and 60% relative humidity. To simulate salt stress, EV, B2, and C4 seedlings were exposed to different concentrations of sodium chloride (NaCl). For root phenotype analyses and isoprene emission measurements, five-day-old seedlings were treated with 0, (mock), 50 (mild stress) or 100 (severe stress) mM NaCl for five days in solid MS medium. To study the primary root growth in soil rather than in an artificial substrate, NE (EV) and IE (B2 and C4) Arabidopsis plants were planted in homemade Rhizoboxes, special pots where root growth can be monitored. Plants were germinated and grown for two weeks in sterile Suremix soil (Michigan Grower Products Inc, Galesburg, MI, USA) and kept in the same controlled growth chambers used for plates.

For RNA-seq, metabolite, and hormone analyses, NE (EV) and IE (B2 and C4) seedlings were grown for 10 days, and then the roots of the seedlings were soaked in 0 or 150 mM NaCl solutions for four h. For RNA-seq analysis, the NE WT was additionally tested, in the same conditions of the other investigated genotypes. This test was intended to verify that the EV did not cause any reprogramming of the transcriptome, confirming previous findings (Zuo et al., 2019; Dani et al., 2022).

### Measurement of isoprene emission from roots

After five days of salt stress treatments, roots were harvested, and 50 mg of the samples were collected in 5 mL glass vials. Roots were incubated for four h at 32°C and a PAR of 300 μmol m^−2^ s^−1^, as described in Miloradovic van Doorn et al. (2020). Isoprene emission was measured in the headspace of the vial with the Fast Isoprene Sensor (FIS, Hills Scientific, Boulder, CO, USA). The headspace gas (3 ml) was injected into air flowing into the instrument at 400 ml min^−1^ by a gastight syringe. The FIS calibration was carried out using a 3.225 ppm isoprene standard (Airgas USA LLC, TX, USA). Isoprene emission was calculated per gram of root fresh weight.

### Metabolites estimation and analysis

#### MEP pathway metabolites

The harvested roots were ground using a tissue-lyser followed by an extraction with an extraction buffer containing acetonitrile: isopropanol: 20 mM ammonium bicarbonate (3:1:1). After centrifugation at 14,000 g for 10 min, the supernatants were collected for analyzing MEP pathway metabolites. A volume of 200 μl supernatant was transferred to glass inserts placed in 2 mL glass vials for LC-MS/MS analysis. MEP pathway metabolites, including 1-deoxy-D-xylulose 5-phosphate (DXP,) 2-C-methyl-D-erythritol 4-phosphate (MEP), methylerythritol cytidyl diphosphate (CDP-ME), 2-C-methyl-D-erythritol-2,4-cyclodiphosphate (MEcDP), 4-hydroxy-3-methyl-butenyl 1-diphosphate (HMBDP), isopentenyl diphosphate (IDP), and dimethylallyl diphosphate (DMADP) were quantified by an Acquity TQD Tandem Quadrupole Mass Spectrometer with an Agilent InfinityLab Poroshell 120 HILIC-Z, column (2.1 x 100 mm, 2.7 μ, Agilent, Santa Clara, CA, USA) following the protocol described by Sahu et al. (2023). Commercial DXP, MEP, CDP-ME, MEcDP, HMBDP, IDP, and DMADP (Logan, UT, USA) were used to develop response curves for calculating the levels of MEP pathway metabolites.

#### Amino acids and organic acids analyses

The extraction of root samples was carried out according to the protocol described by Xu et al. (2021, 2022) with slight modifications. Frozen root samples were ground into a fine powder using a tissue-lyser. The extraction was carried out in chloroform:methanol (3:7) solution for two h at −20°C with vortexing every 30 min. D-[UL-^13^C_6_] fructose 1, 6-bisphosphate and norvaline were added to the sample tubes as internal standards. A volume of 300 μl ice-cold water was added to each tube for extraction of water-soluble metabolites. The Eppendorf tubes were vortexed for 20 s followed by centrifugation at 4,200 g for 10 min. The upper methanol-water phase was separately aliquoted into Eppendorf tubes followed by lyophilization to dryness and stored at −80°C for GC-MS analysis.

For determining the levels of pyruvate, amino acids, and organic acids by GC-MS, we initiated the process by derivatizing the samples with the addition of methoxyamine hydrochloride dissolved in dry pyridine. The mixture was kept at 60°C for 15 min, then cooled for 10 min. Subsequently, it was subjected to silylation by introducing N-tert-bulydimethylasyl-N-methyl-trifluoracetamid with 1% (w/v) tert-bultylmethylchlorisilane, and kept at 60°C overnight, resulting in trimethylsilyl (TBDMS) derivatives. The derivatized samples were then subjected to analysis by an Agilent 7890 GC system (Agilent, Santa Clara, CA, USA) coupled to an Agilent 5975C inert XL Mass Selective Detector (Agilent, Santa Clara, CA, USA) with an autosampler (CTC PAL; Agilent, Santa Clara, CA, USA) following a published protocol (Xu et al., 2022). The characteristic fragment ions used for measuring the metabolites are detailed in Table S1.

#### Hormone contents

Root samples were homogenized using a tissue-lyser, and then extracted in a buffer containing 4:1 methanol:MilliQ water (v/v) added with butylated hydroxytoluene and formic acid. Samples were quantified by Acquity TQD Tandem Quadrupole Mass Spectrometer with an Acquity BEH Amide column (1.7 µm x 2.1 mm x 50 mm) (Acquity Group, Waters, Milford, MA, USA) with an autosampler (2777C, Waters, MA, USA). Salicylic acid (SA), 12-hydroxy jasmonic acid (12OH-JA), jasmonic acid d-5 (JA d-5), abscisic acid d-6 (ABA d-6), 12-hydroxy-jasmonoyl-isoleucine (12OH-JA-Ile), and ^13^C6-indole-3-acetic acid (IAA-^13^C6) were used for developing the standard curves. For quantification of CKs, namely trans-zeatin (tZ), trans-zeatin-riboside (tZR), and isopentenyl-riboside (iPR), root tissues were ground and extracted by 8:2 methanol:MilliQ water (v/v). After centrifugation at 13,000 g for five min, the supernatants were collected and evaporated in a Savant SpeedVac (Thermo-Fisher Scientific, Waltham, MA, USA) at maximum speed for two h. Dried extracts were resuspended in 10% acetonitrile. The samples were run by an Acquity Xevo TQ-XS UPLC/MS/MS (Waters, Milford, MA, USA) and separated with a BEH C18 2.1 x 50 mm column. Caffeine ^13^C_3_ was used as standard. Water and 0.1% formic acid (A) and acetonitrile (B) were used as mobile phase. The mass-spectra acquisition setup included positive mode electrospray ionization mode (ES+), source temperature of 150°C, and desolvation temperature of 400°C. Collision gas and nebulizer gas flow were set to 0.17 ml min^−1^ and 7 bar, respectively. Gas flow for the desolvation and cone was set to 800 and 150 l h^−1^, respectively. Scan time was 100 to 200 amu s^−1^.

#### LC-and GC-MS/MS data analysis

The MassLynx 4.0 and GC/MSD Chemstation (Agilent, Santa Clara, CA, USA) were used to acquire the LC-MS/MS and GC-MS data, respectively. The specific metabolites were identified by their mass to charge (m/z) ratio and retention time using authentic standards. Standard curves were developed using the authentic standards available for each targeted metabolite. Both LC-MS and GC-MS datasets were converted to the MassLynx format, and QuanLynx software was then used to analyze the data, including peak detection and quantification. Absolute quantification of the metabolites was carried out using external standard curves that were normalized with the internal standards.

### Plant phenotype and growth analyses

Plant growth parameters and dry weight (DW) of both roots and shoots were determined after five days of exposure to NaCl solutions. For measuring DW, roots and shoots were separated using a razor blade and transferred to paper bags. The paper bags were oven dried at 65°C for three days and the DW of roots and shoots was then recorded using a digital balance. Root/shoot ratio and percent reduction of root biomass were calculated based on DW. Root phenotype parameters, including primary root (PR) length, lateral root (LR) number, and root growth were measured by analyzing the photographs taken after five days of stress treatments. For determining root gravitropic response, five-d-old NE (EV), and IE (B2 and C4) plants were transferred to new GM plates with fresh MS medium and grown for one additional day. Plates were rotated at 90° immediately after transferring the plants, and the photographs of the roots were recorded at different time points (0, 2, 5, 7, and 24 h) for analyzing the root angles. For Rhizobox analyses, 2-w-old plants were explanted, and pictures were taken for PR length measurements. All root phenotype (plates and soil) parameters were measured using *Fiji* software (Schindelin et al., 2012). All measurements were replicated at least three times.

### Gene expression analyses

#### RNA extraction

Total RNA from frozen root samples was extracted using the RNeasy Mini Kit (Qiagen, Hilden, Germany). The RNA concentration, integrity, and quality were determined with a Qubit RNA Broad-Range Assay Kit (Invitrogen, Waltham, MA, USA) and a Qubit 4 benchtop fluorometer (Invitrogen, Waltham, MA, USA). RNA integrity was further assessed with a 2100 Bioanalyzer (Agilent Technologies, Santa Clara, CA, USA). All samples used for sequencing had an RNA integrity number of at least 8.5 (1-10; low to high quality).

#### Library preparation and RNA-sequencing analysis

mRNA sequencing was performed at the Michigan State University Research Technology Support Facility Genomics Core (https://rtsf.natsci.msu.edu/genomics/). Libraries were prepared using the Illumina Stranded mRNA Library Preparation, Ligation Kit with IDT for Illumina Unique Dual Index adapters following the manufacturer’s recommendations except that half volume reactions were used. Completed libraries were quality checked and quantified using a combination of Qubit dsDNA high sensitivity (HS) and Agilent 4200 Tape Station HS DNA1000 assays. The libraries were pooled in equimolar proportions and the pool quantified using the Invitrogen Collibri Quantification qPCR kit. The pool was combined with other pools of Illumina Stranded RNA libraries prepared by the Genomics Core to make use of a shared S4 lane. This combined pool was likewise quantified using the Invitrogen Collibri Quantification qPCR kit. The combined pool was loaded onto one lane of an Illumina S4 flow cell and sequencing was performed in a 2×150 bp paired end format using a NovaSeq 6000 v1.5, 300 cycle reagent kit. Base calling was done by Illumina Real Time Analysis (RTA) v3.4.4 and output of RTA was demultiplexed and converted to FastQ format with Illumina Bcl2fastq v2.20.0.The paired-end raw sequences obtained from RNA-seq of WT, EV, B2 and C4 grown under both normal (0 mM NaCl) and salt-stress (150 mM NaCl) conditions were checked using FastQC v0.12.1, then trimmed using fastp (Chen et al., 2018) to remove sequences with Q scores < 30. High quality reads were aligned to Athaliana_447_TAIR10.fa reference genome (phytozome-next.jgi.doe.gov/) using STAR 2.7.11a (Dobin et al., 2013). The STAR-output mapped reads were then subjected to featureCounts (Liao et al., 2014), a read summarization program to counts the number of reads mapping to each genomic feature. Estimate variance-mean dependence in read count data and test for differential expressed genes (DEGs) based on negative binomial distribution model was performed using DESeq2 (Love et al., 2014). False discovery rate (FDR < 0.05) and fold change (log_2_ FC ≥ 1.0 or log_2_FC ≤ −1) were used as the minimum cutoffs to determine DEGs among different comparisons.

### Statistical analyses

To statistically determine the effect of isoprene on phenotype (PR length, LR number, FW, DW, and root/shoot ratio), and on the levels of hormones, amino acids, organic acids, and MEP metabolites, data were analyzed by two-way analysis of variance (ANOVA) with post hoc Tukey’s test. PR length reduction, DW reduction of roots, and root bending/curvature were analyzed by two-way ANOVA with post hoc tests. All differences among means were considered statistically significant at *P*< 0.05. Volcano plot analysis was carried out using ‘EnhancedVolcano’ package in RStudio v. 2023.09.0. Protein-protein interaction networks were generated by using multiple proteins search function and *Arabidopsis* as model organism in string database, accessible at (https://string-db.org/).

## Data availability

All the data are available in the main text and in the Supporting Information. All data related to RNA sequencing is currently being submitted and a DOI will be supplied.

## ACKNOWLEDGEMENTS

Authors thank Mass Spec (Dr. Casey Johnny and Dr. Tony Schilmiller) and Genomics Core facilities (Kevin Child) of Michigan State University and their assistance with the experiments. We also thank Cody Keilen (Growth Chamber Facility, Michigan State University) for his assistance and setup of growth chambers and all members of the Sharkey lab for their support. M.B is grateful to “The Company of Biologists” for providing the travelling fellowship award (number DEVTF2210892) (www.biologists.com).

## AUTHOR CONTRIBUTIONS

MB, MGM, and TDS conceived the study and designed the experiments; SMW executed preliminary studies; MB, MGM, and YZ carried out experimental works; MB, MGM, and MA analyzed the data. MB and MGM wrote the original manuscript which was then revised by FL and TDS. TDS, FL, and LDG provided chemicals, reagents, and other research support. All authors revised and approved the manuscript.

**Disclosure statement:** No potential conflict of interest was declared.

## REFERENCES

Arimura G-I (2021) Making sense of the way plants sense herbivores. Trends Plant Sci 26: 288–298

Asensio D, Rapparini F, Peñuelas J (2012) AM fungi root colonization increases the production of essential isoprenoids vs. nonessential isoprenoids especially under drought stress conditions or after jasmonic acid application. Phytochemistry 77: 149–161

Banerjee A, Sharkey TD (2014) Methylerythritol 4-phosphate (MEP) pathway metabolic regulation. Nat Prod Rep 31: 1043–1055

Behnke K, Kaiser A, Zimmer I, Brüggemann N, Janz D, Polle A, Hampp R, Hänsch R, Popko J, Schmitt-Kopplin P, et al (2010) RNAi-mediated suppression of isoprene emission in poplar transiently impacts phenolic metabolism under high temperature and high light intensities: a transcriptomic and metabolomic analysis. Plant Mol Biol 74: 61–75

Bellucci M, Locato V, Sharkey TD, De Gara L, Loreto F (2023) Isoprene emission by plants in polluted environments. J Plant Interact 18: 2266463

Brilli F, Barta C, Fortunati A, Lerdau M, Loreto F, Centritto M (2007) Response of isoprene emission and carbon metabolism to drought in white poplar (Populus alba) saplings. New Phytol 175: 244–254

Chen S, Zhou Y, Chen Y, Gu J (2018) fastp: an ultra-fast all-in-one FASTQ preprocessor. Bioinformatics 34: i884–i890

Cinege G, Louis S, Hänsch R, Schnitzler J-P (2009) Regulation of isoprene synthase promoter by environmental and internal factors. Plant Mol Biol 69: 593–604

Corwin DL (2021) Climate change impacts on soil salinity in agricultural areas. Eur J Soil Sci 72: 842–862

Dani KGS, Loreto F (2022) Plant volatiles as regulators of hormone homeostasis. New Phytol 234: 804–812

Dani KGS, Pollastri S, Pinosio S, Reichelt M, Sharkey TD, Schnitzler J-P, Loreto F (2022) Isoprene enhances leaf cytokinin metabolism and induces early senescence. New Phytol 234: 961–974

Dani KGS, Torzillo G, Michelozzi M, Baraldi R, Loreto F (2020) Isoprene emission in darkness by a facultative heterotrophic green alga. Front Plant Sci 11: 598786

Dobin A, Davis CA, Schlesinger F, Drenkow J, Zaleski C, Jha S, Batut P, Chaisson M, Gingeras TR (2013) STAR: ultrafast universal RNA-seq aligner. Bioinformatics 29: 15–21

Frank L, Wenig M, Ghirardo A, van der Krol A, Vlot AC, Schnitzler J-P, Rosenkranz M (2021) Isoprene and β-caryophyllene confer plant resistance via different plant internal signalling pathways. Plant Cell Environ 44: 1151–1164

Galvan-Ampudia CS, Testerink C (2011) Salt stress signals shape the plant root. Curr Opin Plant Biol 14: 296–302

Ghirardo A, Gutknecht J, Zimmer I, Brüggemann N, Schnitzler J-P (2011) Biogenic volatile organic compound and respiratory CO2 emissions after 13C-labeling: online tracing of C translocation dynamics in poplar plants. PLoS One 6: e17393

Guenther A, Karl T, Harley P, Wiedinmyer C, Palmer PI, Geron C (2006) Estimates of global terrestrial isoprene emissions using MEGAN (Model of Emissions of Gases and Aerosols from Nature). Atmos Chem Phys 6: 3181–3210

Harvey CM, Sharkey TD (2016) Exogenous isoprene modulates gene expression in unstressed Arabidopsis thaliana plants. Plant Cell Environ 39: 1251–1263

Huque AKMM, So W, Noh M, You MK, Shin JS (2021) Overexpression of AtBBD1, Arabidopsis bifunctional nuclease, confers drought tolerance by enhancing the expression of regulatory genes in ABA-Mediated drought stress signaling. Int J Mol Sci 22: 2936

Dello Ioio R, Nakamura K, Moubayidin L, Perilli S, Taniguchi M, Morita MT, Aoyama T, Costantino P, Sabatini S (2008) A genetic framework for the control of cell division and differentiation in the root meristem. Science (80-) 322: 1380–1384

Jardine K, Monson R, Abrell L, Saleska S, Arneth A, Jardine AB, Ishida Y, Yañez-Serrano A, Artaxo P, Karl T, et al (2012) Within-plant isoprene oxidation confirmed by direct emissions of oxidation products methyl vinyl ketone and methacrolein. Glob Chang Biol 18: 973–984

Julkowska MM, Koevoets IT, Mol S, Hoefsloot H, Feron R, Tester MA, Keurentjes JJB, Korte A, Haring MA, de Boer G-J, et al (2017) Genetic components of root architecture remodeling in response to salt stress. Plant Cell 29: 3198–3213

Karlova R, Boer D, Hayes S, Testerink C (2021) Root plasticity under abiotic stress. Plant Physiol 187: 1057–1070

Kigathi RN, Weisser WW, Reichelt M, Gershenzon J, Unsicker SB (2019) Plant volatile emission depends on the species composition of the neighboring plant community. BMC Plant Biol 19: 58

Lantz AT, Solomon C, Gog L, McClain AM, Weraduwage SM, Cruz JA, Sharkey TD (2019) Isoprene suppression by CO2 is not due to Triose Phosphate Utilization (TPU) limitation. Front For Glob Chang.

Liao Y, Smyth GK, Shi W (2014) featureCounts: an efficient general purpose program for assigning sequence reads to genomic features. Bioinformatics 30: 923–930

Loivamäki M, Gilmer F, Fischbach RJ, Sörgel C, Bachl A, Walter A, Schnitzler J-P (2007) Arabidopsis, a model to study biological functions of isoprene emission? Plant Physiol 144: 1066–1078

Loreto F, Delfine S (2000) Emission of isoprene from salt-stressed Eucalyptus globulus leaves. Plant Physiol 123: 1605–1610

Loreto F, Mannozzi M, Maris C, Nascetti P, Ferranti F, Pasqualini S (2001) Ozone quenching properties of isoprene and its antioxidant role in leaves. Plant Physiol 126: 993– 1000

Loreto F, Schnitzler J-P (2010) Abiotic stresses and induced BVOCs. Trends Plant Sci 15: 154–166

Love MI, Huber W, Anders S (2014) Moderated estimation of fold change and dispersion for RNA-seq data with DESeq2. Genome Biol 15: 550

Miloradovic van Doorn M, Merl-Pham J, Ghirardo A, Fink S, Polle A, Schnitzler J-P, Rosenkranz M (2020) Root isoprene formation alters lateral root development. Plant Cell Environ 43: 2207–2223

Minhas PS, Ramos TB, Ben-Gal A, Pereira LS (2020) Coping with salinity in irrigated agriculture: Crop evapotranspiration and water management issues. Agric Water Manag 227: 105832

Monson RK, Weraduwage SM, Rosenkranz M, Schnitzler J-P, Sharkey TD (2021) Leaf isoprene emission as a trait that mediates the growth-defense tradeoff in the face of climate stress. Oecologia 197: 885–902

Monson RK, Winkler B, Rosenstiel TN, Block K, Merl-Pham J, Strauss SH, Ault K, Maxfield J, Moore DJP, Trahan NA, et al (2020) High productivity in hybrid-poplar plantations without isoprene emission to the atmosphere. Proc Natl Acad Sci U S A 117: 1596–1605

Morfopoulos C, Sperlich D, Peñuelas J, Filella I, Llusià J, Medlyn BE, Niinemets Ü, Possell M, Sun Z, Prentice IC (2014) A model of plant isoprene emission based on available reducing power captures responses to atmospheric CO2. New Phytol 203: 125–139

Moubayidin L, Di Mambro R, Sabatini S (2009) Cytokinin-auxin crosstalk. Trends Plant Sci 14: 557–562

Pollastri S, Baccelli I, Loreto F (2021) Isoprene: an antioxidant itself or a molecule with multiple regulatory functions in plants? Antioxidants 10: 684

Pollastri S, Jorba I, Hawkins TJ, Llusià J, Michelozzi M, Navajas D, Peñuelas J, Hussey PJ, Knight MR, Loreto F (2019) Leaves of isoprene-emitting tobacco plants maintain PSII stability at high temperatures. New Phytol 223: 1307–1318

Rai VK (2002) Role of amino acids in plant responses to stresses. Biol Plant 45: 481–487

Sahu A, Mostofa MG, Weraduwage SM, Sharkey TD (2023) Hydroxymethylbutenyl diphosphate accumulation reveals MEP pathway regulation for high CO2-induced suppression of isoprene emission. Proc Natl Acad Sci 120: e2309536120

Saxena A, Mishra S, Ray S, Raghuwanshi R, Singh HB (2020) Differential reprogramming of defense network in Capsicum annum L. plants against Colletotrichum truncatum infection by phyllospheric and rhizospheric Trichoderma strains. J Plant Growth Regul 39: 751–763

Schindelin J, Arganda-Carreras I, Frise E, Kaynig V, Longair M, Pietzsch T, Preibisch S, Rueden C, Saalfeld S, Schmid B, et al (2012) Fiji: an open-source platform for biological-image analysis. Nat Methods 9: 676–682

Sharkey TD, Gray DW, Pell HK, Breneman SR, Topper L (2013) Isoprene synthase genes form a monophyletic clade of acyclic terpene synthases in the TPS-B terpene synthase family. Evolution (N Y) 67: 1026–1040

Sharkey TD, Monson RK (2017) Isoprene research - 60 years later, the biology is still enigmatic. Plant Cell Environ 40: 1671–1678

Sharkey TD, Yeh S (2001) Isoprene emission from plants. Annu Rev Plant Physiol Plant Mol Biol 52: 407–436

Tattini M, Velikova V, Vickers C, Brunetti C, Di Ferdinando M, Trivellini A, Fineschi S, Agati G, Ferrini F, Loreto F (2014) Isoprene production in transgenic tobacco alters isoprenoid, non-structural carbohydrate and phenylpropanoid metabolism, and protects photosynthesis from drought stress. Plant Cell Environ 37: 1950–1964

Velikova V, Loreto F (2005) On the relationship between isoprene emission and thermotolerance in Phragmites australis leaves exposed to high temperatures and during the recovery from a heat stress. Plant Cell Environ 28: 318–327

Velikova V, Pinelli P, Pasqualini S, Reale L, Ferranti F, Loreto F (2005) Isoprene decreases the concentration of nitric oxide in leaves exposed to elevated ozone. New Phytol 166: 419– 425

Vickers C, Possell M, Cojocariu C, Velikova V, Laothawornkitkul J, Ryan A, Mullineaux P, Hewitt CN (2009) Isoprene synthase protects transgenic tobacco plants from oxidative stress. Plant Cell Environ 32: 520–531

Vlot AC, Liu P-P, Cameron RK, Park S-W, Yang Y, Kumar D, Zhou F, Padukkavidana T, Gustafsson C, Pichersky E, et al (2008) Identification of likely orthologs of tobacco salicylic acid-binding protein 2 and their role in systemic acquired resistance in Arabidopsis thaliana. Plant J 56: 445–456

Weise SE, Li Z, Sutter AE, Corrion A, Banerjee A, Sharkey TD (2013) Measuring dimethylallyl diphosphate available for isoprene synthesis. Anal Biochem 435: 27–34

Weraduwage SM, Whitten D, Kulke M, Sahu A, Vermaas JV, Sharkey TD (2023) The isoprene-responsive phosphoproteome provides new insights into the putative signalling pathways and novel roles of isoprene. Plant Cell Environ 10.1111/pce.14776

Xu J, Trainotti L, Li M, Varotto C (2020) Overexpression of isoprene synthase affects ABA-and drought-related gene expression and enhances tolerance to abiotic stress. Int J Mol Sci 21: 4276

Xu Y, Fu X, Sharkey TD, Shachar-Hill Y, Walker BJ (2021) The metabolic origins of non-photorespiratory CO2 release during photosynthesis: a metabolic flux analysis. Plant Physiol 186: 297–314

Xu Y, Wieloch T, Kaste JAM, Shachar-Hill Y, Sharkey TD (2022) Reimport of carbon from cytosolic and vacuolar sugar pools into the Calvin-Benson cycle explains photosynthesis labeling anomalies. Proc Natl Acad Sci U S A 119: e2121531119

Yang Y, Xu R, Ma C-J, Vlot AC, Klessig DF, Pichersky E (2008) Inactive methyl indole-3-acetic acid ester can be hydrolyzed and activated by several esterases belonging to the AtMES esterase family of Arabidopsis. Plant Physiol 147: 1034–1045

Zhou F, Pichersky E (2020) The complete functional characterisation of the terpene synthase family in tomato. New Phytol 226: 1341–1360

Zou Y, Zhang Y, Testerink C (2022) Root dynamic growth strategies in response to salinity. Plant Cell Environ 45: 695–704

Zuo Z, Weraduwage SM, Lantz AT, Sanchez LM, Weise SE, Wang J, Childs KL, Sharkey TD (2019) Isoprene acts as a signaling molecule in gene networks important for stress responses and plant growth. Plant Physiol 180: 124–152

